# The translocation mechanism of calcitriol through *Helicobacter pylori* lipid membrane and influence on water permeability

**DOI:** 10.1101/2023.03.22.533895

**Authors:** Zanxia Cao, Liling Zhao, Mingcui Chen, Lei Liu

**Author notes:** Correspondence (Z.C.), (Z.C.), (L.L.), (Z.C.). (LL.Z.); (MC.C).

## Abstract

*Helicobacter pylori* exhibits a unique membrane lipid composition, including dimyristoyl phosphatidylethanolamine (DMPE) and cholesterol, unlike other Gram-negative bacteria. Calcitriol has antimicrobial activity against *H. pylori*, but cholesterol enhances antibiotics resistance in *H. pylori*. This study explored the changes in membrane structure and the molecular mechanisms of cholesterol/calcitriol translocation using well-tempered metadynamics (WT-MetaD) simulations and microsecond conventional molecular dynamics simulations. Our results showed that the average area per lipid and sterol tilt angles were slightly lower, while D_P-P_, D_CG-CG_, D_AC-AC_, and S_CD_ were higher in cholesterol membrane systems than in calcitriol membrane systems. Cholesterol membrane systems were more ordered than calcitriol-containing membranes. Calcitriol facilitated water transport across the membrane, while cholesterol had the opposite effect. The differing effects might result from the tail 25-hydroxyl group and a wider range of orientations of calcitriol in the DMPE/ dimyristoyl phosphatidylglycerol (DMPG) (3:1) membrane. Calcitriol moves across the bilayer center without changing its orientation along the membrane Z-axis, becomes parallel to the membrane surface at the membrane-water interface, and then rotates approximately 90º in this interface. The translocation mechanism of calcitriol is quite different from the flip-flop of cholesterol. Moreover, calcitriol crossed from one layer to another more easily than cholesterol, causing successive perturbations to the hydrophobic core and increasing water permeation. These results improve our understanding of the relationship between cholesterol/ calcitriol concentrations and the lipid bilayer structure and the role of lipid composition in water permeation.

## 1. Introduction

*Helicobacter pylori* is a highly prevalent Gram-negative bacterium that causes peptic ulcers and gastric cancer. This bacterium exhibits a unique membrane lipid composition. Dimyristoyl phosphatidylethanolamine (DMPE) with a myristic acid (C14:0) as the saturated fatty acid side chain is one of the most abundant lipid components in *H. pylori* membranes, in contrast to other Gram-negative bacteria. Moreover, *H. pylori* incorporates exogenous cholesterol into the cell membrane and uses cholesterol to acquire resistance to antibiotics, LL-37[1] and lipophilic compounds[2]. DMPE strongly and selectively binds to cholesterol [3, 4].

Cholesterol maintains membrane fluidity, decreases the average area per lipid (APL), and increases membrane thickness[5]. These characteristics reduce water permeability. In line with these findings, molecular dynamics (MD) simulations indicated that the presence of cholesterol in the 1-palmitoyl-2-oleoyl-sn-glycero-3-phosphatidylcholine (POPC) bilayer[6], 1,2-dipalmitoyl-sn-glycero-3-phosphatidylcholine (DPPC) bilayer[7], phosphatidylglycerol bilayer[8] and dioleylphosphatidylcholine (DOPC)/ sphingomyelin bilayer[9] significantly reduced water permeability, demonstrating that reduced water permeability was caused by the increased organization of lipids in the presence of cholesterol.

Calcitriol (1,25-dihydroxyvitamin D) is the active form of vitamin D, and cholesterol is the precursor of vitamin D (cholecalciferol). Calcitriol and cholesterol have 74 atoms (Figure 1C). Cholesterol is a 27-carbon compound with one hydroxyl group, a central sterol nucleus containing four hydrocarbon rings, and a hydrocarbon tail. Calcitriol is a 27-carbon compound with three hydroxyl groups, a central sterol nucleus, and a hydrocarbon tail. The hydroxyl groups are polar, and the central sterol and hydrocarbon tails are nonpolar. Calcitriol has antimicrobial activity against *H. pylori* [4, 10, 11], and 100 μM calcitriol kills this bacterium (108.5 CFU/mL) within 2 h. In addition, clinical results showed that serum 25-hydroxyvitamin D levels lower than 20 ng/mL might affect the ability of quadruple therapy to kill *H. pylori* [12], and higher levels were associated with effective killing [13]. Nonetheless, little is known about the behavior of calcitriol in the *H. pylori* membrane bilayer.

**Figure 1.**
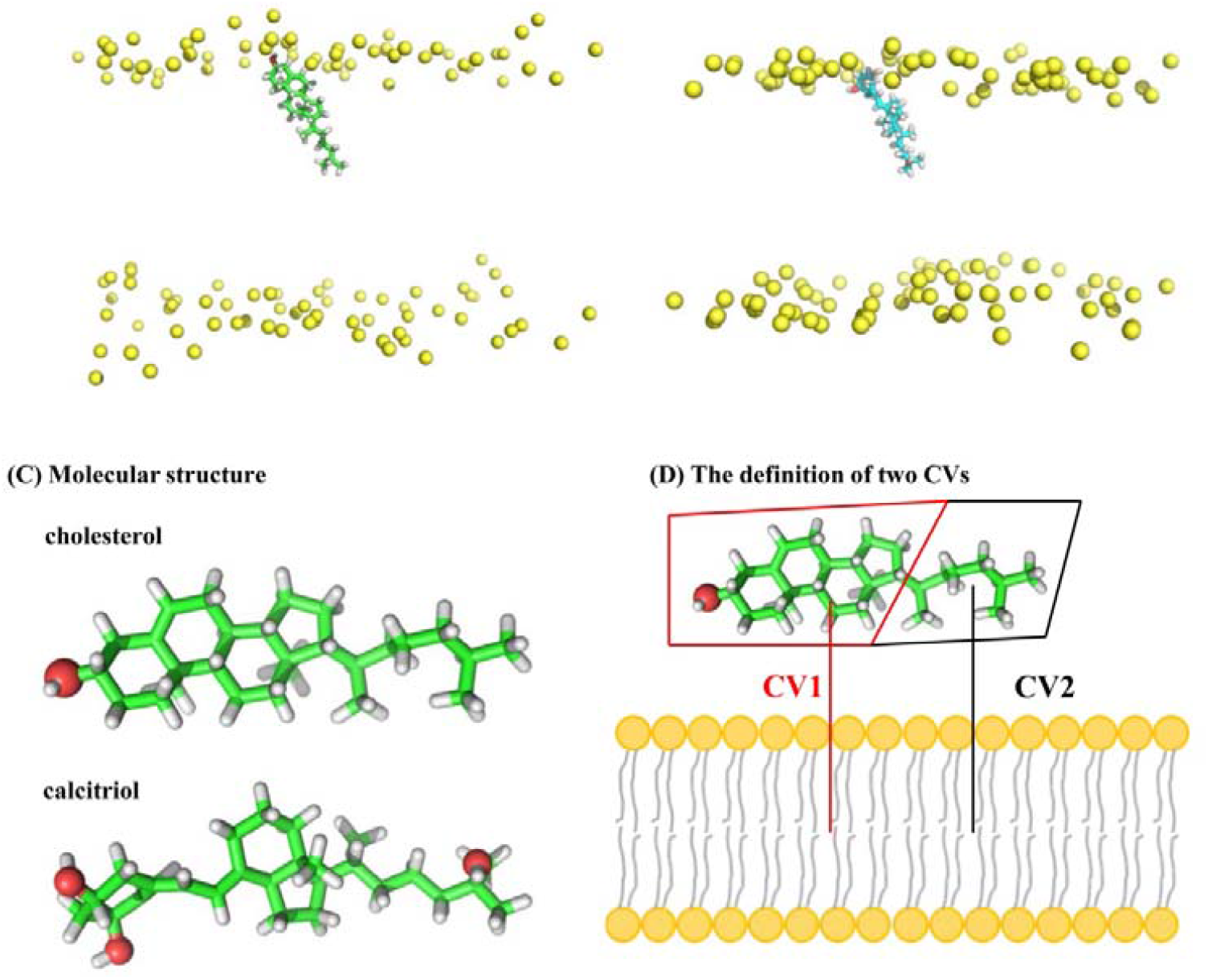
Initial conformation of (A) the cholesterol-DMPE/DMPG (3:1) system and (B) the calcitriol-DMPE/DMPG (3:1) system. (C-D) Chemical structure of cholesterol and calcitriol and definition of two collective variables. Hydrophilic heads, cholesterol/calcitriol molecules, and oxygen atoms of hydroxyl groups are shown in yellow, green, and red, respectively.

Based on the above considerations, this study investigated the changes in membrane structure and the molecular mechanisms of cholesterol/calcitriol translocation using well-tempered metadynamics (WT-MetaD) simulations and microsecond conventional molecular dynamics simulations. The DMPE/DMPG (3:1) lipid bilayer was used as a model of the *H. pylori* membrane. Changes in membrane structural characteristics using different concentrations of cholesterol and calcitriol were compared. These data provide a basis to improve the design of anti-*H. pylori* agents.

## 2. METHODS

### 2.1 WT-MetaD simulations in cholesterol- and calcitriol-containing membrane systems

The DMPE/DMPG (3:1) lipid bilayer was used as a model to analyze the lipid characteristics of the *H. pylori* membrane. The membranes contained 96 DMPE lipids and 32 DMPG lipids. The structure and topology of the pure lipid bilayer were created using CHARMM-GUI *Membrane Builder* (http://charmm-gui.org/)[14, 15]. To obtain an equilibrated system, a pure DMPE/DMPG (96:32) lipid bilayer was simulated for 300 ns, and the last snapshot was used as the membrane system. Each simulation contained one cholesterol or calcitriol molecule, which was placed approximately 1.2 nm from the bilayer center in the vertical orientation. The initial conformation of these simulations is shown in Figures 1A and B.

All WT-MetaD simulations[16] were performed using GROMACS 2018 software package[17]. This work was done using the open-source, community-developed PLUMED library version 2.5[18]. We employed two collective variables (CVs): (1) the Z distance between the center of mass of the indene and the membrane and (2) the Z distance between the alkyl and the membrane. The molecular structures are shown in Figure 1C, and the definitions of CVs are shown in Figure 1D. The Chemistry at Harvard Molecular Mechanics (CHARMM) force field[19] was adopted to simulate the DMPE/DMPG (96:32) lipid bilayer and cholesterol. The force field parameters of calcitriol were obtained using CGenFF[20] (https://cgenff.umaryland.edu/). We applied periodic boundary condition in a rectangle (5.25 nm × 5.25 nm × 16 nm for DMPE/DMPG [96:32]). The systems were solvated using the TIP3P water model and neutralized by adding 32 Na^+^ cations to the simulation box. After extensive energy minimization, 5 ns equilibration simulations in the NVT and NPT ensembles were performed. The temperature was controlled at 333 K using the Nose-Hoover algorithm[21] with a tau-t of 1 ps. The pressure was maintained at 1 atm using the semi-isotropic Parrinello-Rahman method[22] with a tau-p of 5 ps and a compressibility of 4.5 × 10 –5 bar –1. Long-range electrostatic interactions with a grid spacing of 0.12 nm were treated using the Particle Mesh Ewald method[23]. The LJ interaction with a cut-off of 1 nm was calculated using the force-based switching function. The bond length was constrained using the LINCS algorithm[24]. The initial Gaussian height was 0.8 kJ/mol, and the width was 0.2 nm for both CVs. The bias factor was set to 10. The total time of the simulation was 1000 ns.

### 2.2 CMD simulations

We performed CMD simulations using different numbers of cholesterol and calcitriol molecules in DMPE/DMPG (3:1) membranes (Table 1). The distance between the bilayer center and the cholesterol/calcitriol molecule in the global minimum energy conformations of WT-MetaD simulations was set as a reference to construct membrane systems containing multiple cholesterol or calcitriol molecules. All simulated systems were equilibrated for 800 ns, and subsequent 200 ns simulations were used for analysis.

**Table 1.**
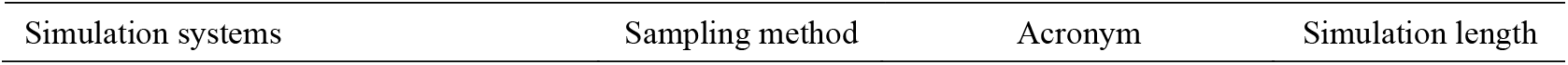
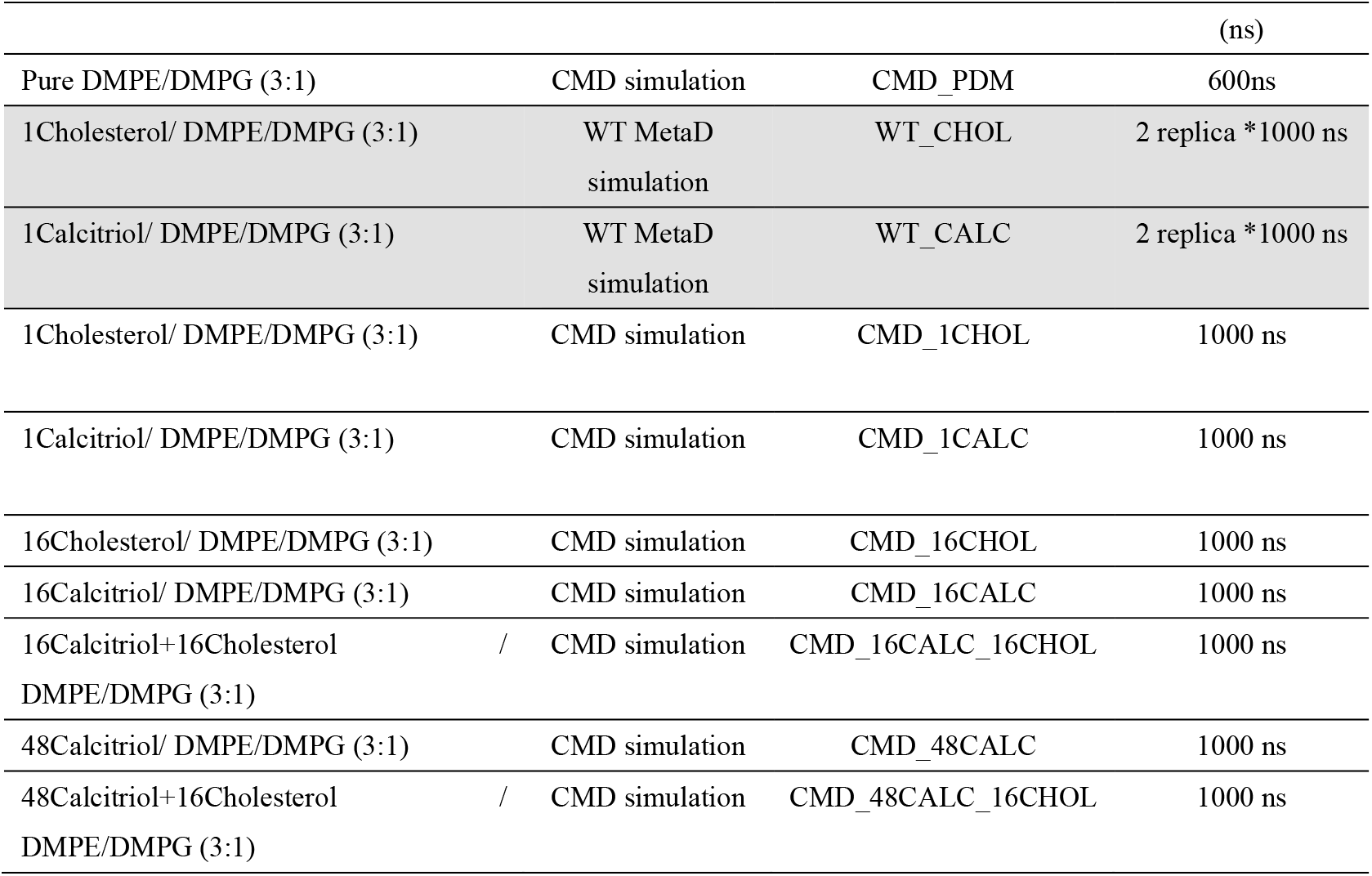
Summary of molecular dynamics simulations.

Data description

The input file of simulation https://github.com/zxcao/translocation-mechanism-of-calcitriol

## 3. RESULTS

### 3.1 Two-dimensional free energy landscapes (FELs) from WT-MetaD simulations

The two-dimensional free energy (ΔG) for the translocation of cholesterol or calcitriol into the DMPE/DMPG (3:1) bilayer along the two CVs is shown in Figure 2. The FELs in membranes containing one molecule of cholesterol or calcitriol are shown in Figure 2. A major difference between the membrane systems was the magnitude of the global free energy minimum. The energy minimum of the cholesterol-DMPE/DMPG (3:1) system was located at CV1=1.4 nm and CV2=0.0 nm within -80.0 kJ/mol. The energy minimum of the calcitriol-DMPE/DMPG (3:1) system was located at CV1=1.0 nm and CV2=0.2 nm within -53.0 kJ/mol. Global minimum energy conformations are shown in Figure 3. Cholesterol and calcitriol were inserted into the bilayer membrane and oriented along the bilayer Z-axis.

**Figure 2.**
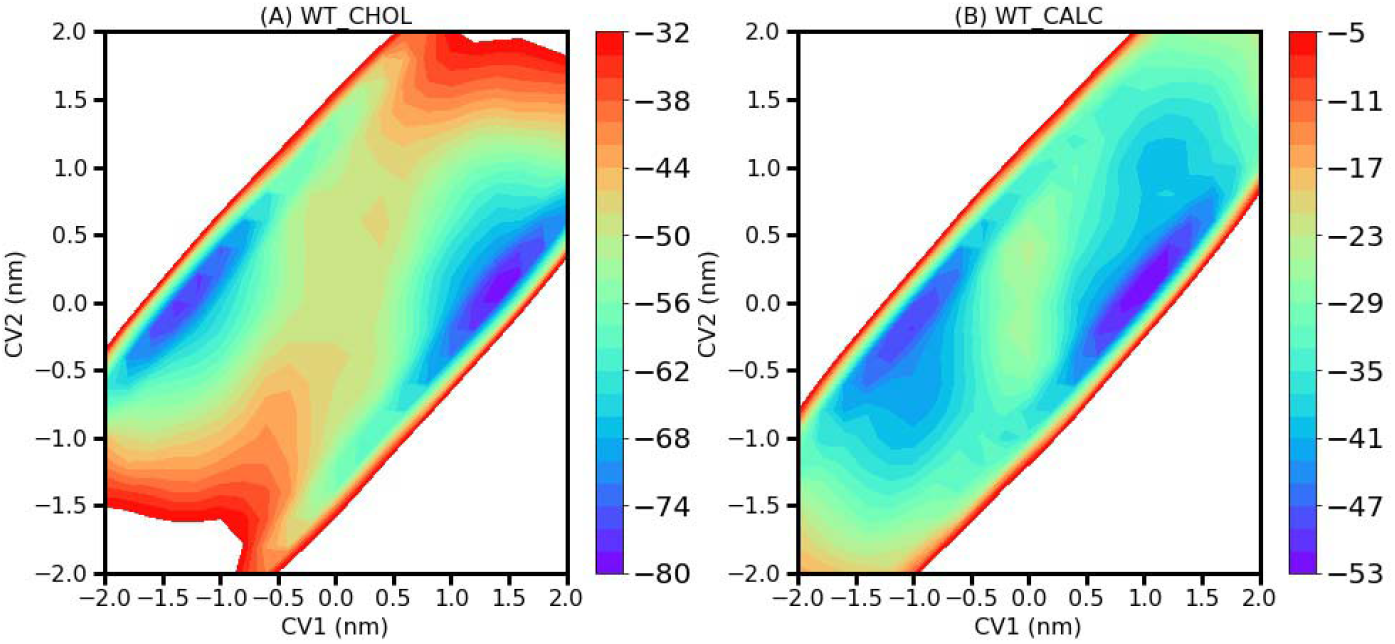
Two-dimensional free energy landscapes of (A) the cholesterol-DMPE/DMPG (3:1) system and (B) the calcitriol-DMPE/DMPG (3:1) system along two collective variables. The free energy value is set to zero in the water phase.

**Figure 3.**
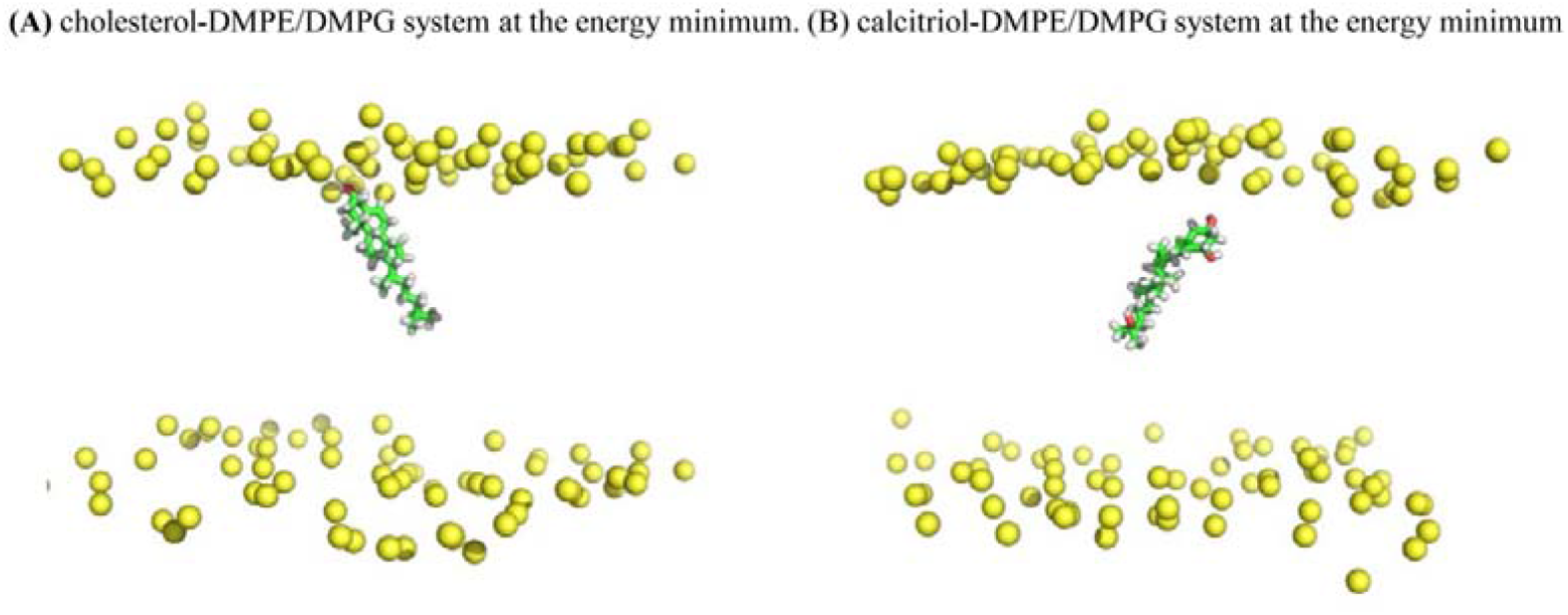
The global minimum energy conformations of (A) the cholesterol-DMPE/DMPG (3:1) system and (B) the calcitriol-DMPE/DMPG (3:1) system. Hydrophilic heads, cholesterol/calcitriol molecules, and oxygen atoms of hydroxyl groups are shown in yellow, green, and red, respectively.

The local energy minimum was narrow for the cholesterol-DMPE/DMPG (3:1) bilayer system, and cholesterol was positioned vertically into the membrane. CV1 and CV2 were 1.0-1.8 nm and -0.4–0.4 nm away from the bilayer center, respectively. Conversely, the local energy minimum was wide for the calcitriol-DMPE/DMPG (3:1) bilayer system, indicating that calcitriol was positioned vertically and horizontally into the membrane. CV1 and CV2 were 0.5-1.5 nm and -0.3–1.5 nm from the bilayer center, respectively. Calcitriol had a wider range of motion than cholesterol in this membrane system.

### 3.2 Structural and dynamic changes in membranes based on CMD simulations

The *H. pylori* membrane is the primary target for most antibacterial compounds. The deuterium order parameters (*S*_CD_) describe the molecular motions of acyl chains and are an important indicator of the membrane stability. The *S*_CD_ value is positively correlated with the order of the hydrophobic core. Therefore, the *S*_CD_ of acyl chains was calculated using the Membrainy tool[25] in pure DMPE/DMPG (3:1) lipid bilayers and membrane systems with a varying number of cholesterol and calcitriol molecules (Figure 4A). The order of acyl chains for the DMPE/DMPG/cholesterol/calcitriol systems increased with increasing sterol concentration (Figure 4A). Therein, the hydroxy groups of cholesterol and calcitriol were oriented close to the glycerol and phosphate head groups of the DMPE/DMPG lipid bilayer, which decreased the mobility of the hydrocarbon chain in this region. The lipid order increased as follows: CMD_PDM < CMD_1CALC<CMD_1CHOL < CMD_16CALC < CMD_48CALC < CMD_16CHOL < CMD_16CALC_16CHOL <

**Figure 4.**
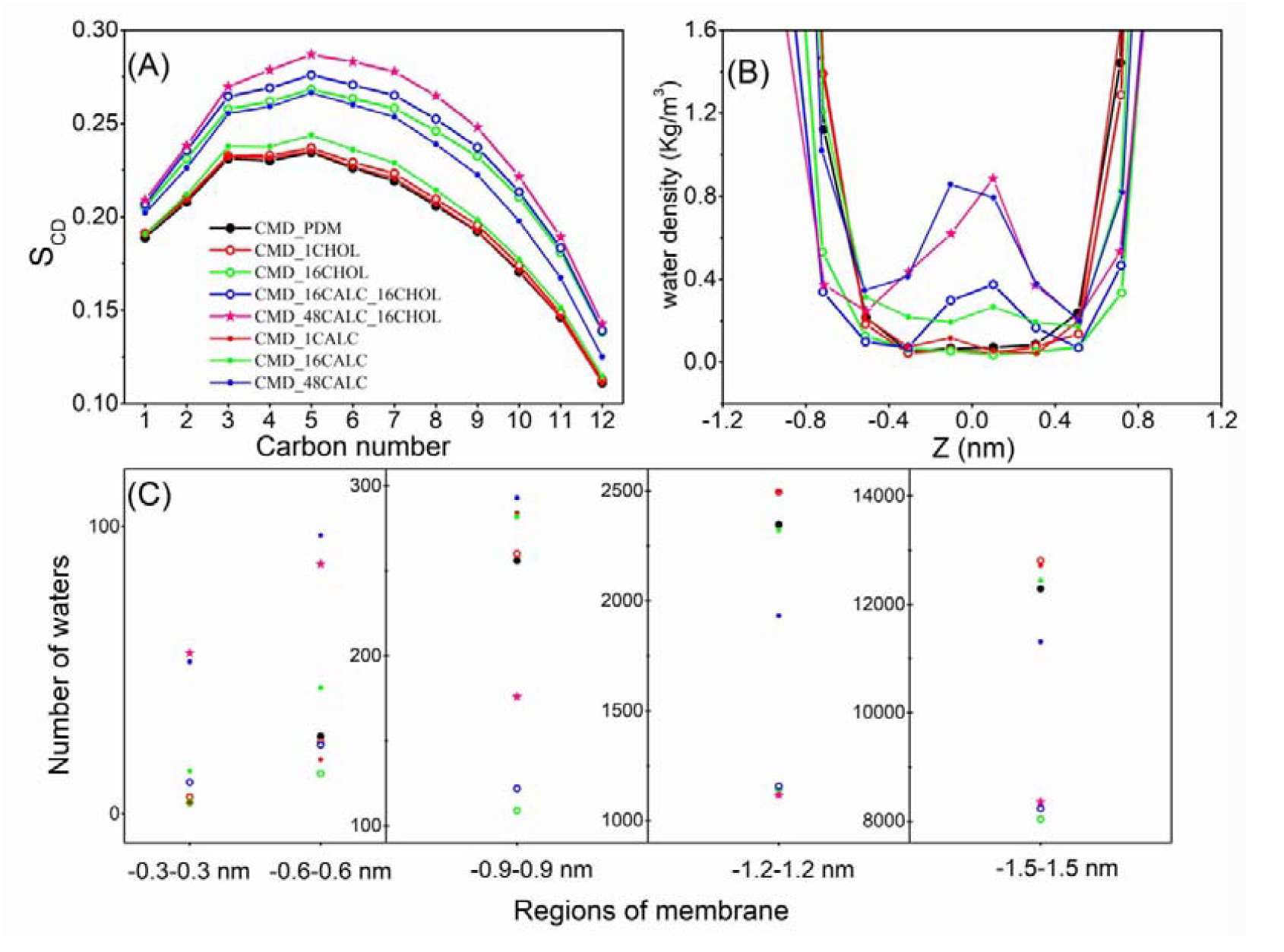
(A). Deuterium order parameters (S_CD_) for the lipid acyl chains. (B). Mass density of water in conventional molecular dynamics simulations using a varying number of cholesterol and calcitriol molecules. (C). Number of water molecules in different regions of the lipid bilayer.

CMD_48CALC_16CHOL. The simulations indicated that the organization of calcitriol was significantly lower than that of cholesterol in this membrane system. Moreover, DMPE/DMPG (3:1) lipid bilayers containing 16 cholesterol molecules were more ordered than bilayers containing 16 calcitriol molecules.

The following membrane properties were also analyzed: membrane area (X = Y; thus, only the X-axis is shown), membrane thickness between phosphate groups (D_P-P_), membrane thickness between carbonyl groups (D_CG-CG_), membrane thickness between the first aliphatic carbons (D_AC-AC_), and sterol tilt angle. The APL was calculated by dividing the membrane area by the total number of lipids in each leaflet. The sterol tilt angle was calculated as the angle between the vector of sterols (from C17 to C3) and the membrane Z-axis. The average APL and sterol tilt angles were slightly lower, while D_P-P_, D_CG-CG_, D_AC-AC_, and S_CD_ were higher in cholesterol-containing membranes than in membrane systems with the corresponding number of calcitriol molecules (Table 2). These results indicate that calcitriol-containing membranes are more disordered than cholesterol-containing membranes. Looser packing of lipids in the former facilitates the transmembrane diffusion of lipids, proteins, and other molecules.

**Table 2.** Membrane thickness between phosphate groups (D_P-P_), membrane thickness between carbonyl groups (D_CG-CG_), membrane thickness between the first aliphatic carbons (D_AC-AC_), X dimension, and average tilt angle were calculated for all systems using conventional molecular dynamics simulations.

| Simulation | $D_{P-P}$ (nm) | $D_{CG-CG}$ (nm) | $D_{AC-AC}$ (nm) | X dimension (nm) | Cholesterol tilt angle | Calcitriol tilt angle |
| --- | --- | --- | --- | --- | --- | --- |
| CMD_PDM | 3.65±0.07 | 2.77±0.06 | 2.58±0.06 | 6.19±0.07 |  |  |
| CMD_1CHOL | 3.67±0.07 | 2.79±0.06 | 2.59±0.06 | 6.22±0.07 | 23±13 |  |
| CMD_1CALC | 3.65±0.07 | 2.78±0.06 | 2.58±0.06 | 6.19±0.07 |  | 32±18 |
| CMD_16CHOL | 3.83±0.06 | 2.95±0.06 | 2.75±0.06 | 6.27±0.06 | 20±11 |  |
| CMD_16CALC | 3.68±0.06 | 2.81±0.06 | 2.61±0.06 | 6.40±0.06 |  | 30±17 |
| CMD_16CALC_16CHOL | 3.85±0.06 | 2.98±0.06 | 2.78±0.06 | 6.50±0.07 | 20±11 | 27±15 |
| CMD_48CALC | 3.74±0.06 | 2.88±0.05 | 2.68±0.05 | 6.80±0.07 |  | 28±16 |
| CMD_48CALC_16CHOL | 3.88±0.06 | 3.01±0.05 | 2.81±0.05 | 6.92±0.06 | 20±11 | 26±15 |

The water leakage effect induced by molecules was measured by calculating the Z-dependent water mass density (Figure 4B) and by counting the number of water molecules (Figure 4C) in different regions of the membrane. The membrane was centered at Z = 0. The mass density of water in membranes with 16 or 48 calcitriol molecules was higher than in membranes without calcitriol. The bilayer core—a 0.6 nm region on the mid-plane of the bilayer—contained the largest number of water molecules in simulations with 48 calcitriol molecules regardless of the presence of cholesterol. In simulation CMD_16CHOL, all regions contained the least number of water molecules. Acyl chains were more ordered than the pure DMPE/DMPG membrane in the presence of calcitriol but could carry water molecules into the center of the membrane.

### 3.3 Interactions between cholesterol/calcitriol and membranes

The binding free energies of the single cholesterol-DMPE/DMPG (3:1) bilayer and single calcitriol-DMPE/DMPG (3:1) bilayer from the last 200 ns in simulations CMD_1CHOL and CMD_1CALC (Table 3) were calculated using g_mmpbsa[26]. The total binding free energy was obtained using equation (1) and the molecular mechanics Poisson Boltzmann surface area method[26].

**Table 3.** Binding free energy (kJ/mol) between a single cholesterol molecule or a single calcitriol molecule and the DMPE/DMPG (3:1) lipid bilayer during the last 200 ns trajectories in simulations CMD_1CHOL and CMD_1CALC.

| | $\Delta G_{\text{bind}}$ | $\Delta E_{\text{ele}}$ | $\Delta E_{\text{vdw}}$ | $\Delta G_{\text{pol}}$ | $\Delta G_{\text{nonpol}}$ |
| --- | --- | --- | --- | --- | --- |
| CMD_1CHOL | $-212 \pm 14$ | $-12 \pm 11$ | $-198 \pm 13$ | $51 \pm 23$ | $-52 \pm 3$ |
| CMD_1CALC | $-161 \pm 26$ | $-29 \pm 13$ | $-194 \pm 15$ | $85 \pm 27$ | $-22 \pm 5$ |

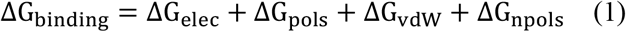

Where ΔG_elec_, ΔG_pols_, ΔG_vdw_, and ΔG_npols_ are the electrostatic energy, polar solvation energy, van der Waals energy, and nonpolar solvation energy, respectively. The binding free energies of the cholesterol-DMPE/DMPG (3:1) bilayer and calcitriol-DMPE/DMPG (3:1) bilayer were -212 and -161 kJ/mol, respectively. The binding free energy was decomposed into individual energies to further explore these results. The sum of *van der Waals* interactions and nonpolar solvation energies of these bilayers was -250 and -216 kJ/mol, respectively. However, the electrostatic contribution (the sum of the electrostatic interaction energy and polar solvation energy) of these bilayers was 39 and 56 kJ/mol, respectively, indicating that the contribution of nonpolar interactions was greater in the cholesterol-DMPE/DMPG (3:1) bilayer system.

The number of hydrogen bonds and contacts between cholesterol/calcitriol and the DMPE/DMPG (3:1) bilayer was calculated from the last 200 ns of the CMD trajectories (Tables 4 and 5). The hydroxyl groups of cholesterol and calcitriol can be used as acceptors or donors to form hydrogen bonds with the bilayer membrane or water. In CMD_1CHOL, the average number of hydrogen bonds between a cholesterol molecule and the membrane was approximately 0.5, and the average number of hydrogen bonds between a cholesterol molecule and water was approximately 1.4. The total number of hydrogen bonds increased linearly with the number of cholesterol molecules, while the number of hydrogen bonds between a cholesterol molecule and the membrane remained unchanged. The presence of calcitriol did not significantly affect the number of hydrogen bonds between cholesterol and the membrane/water. In CMD_1CALC, the average number of hydrogen bonds between a calcitriol molecule and the membrane was approximately 1.0, and the average number of hydrogen bonds between a calcitriol molecule and water was approximately 1.8. The number of hydrogen bonds between single calcitriol and the membrane decreased gradually with the number of calcitriol molecules. Looser packing of lipids in calcitriol-containing membranes inhibited the interaction between calcitriol and the DMPE/DMPG membrane. The presence of cholesterol did not significantly affect the number of hydrogen bonds between calcitriol and the membrane/water. The number of hydrogen bonds between calcitriol and water was higher than that between cholesterol and water.

**Table 4.** Number of hydrogen bonds in cholesterol-membrane, cholesterol-water, calcitriol-membrane, and calcitriol-water interactions during conventional molecular dynamics simulations.

| Simulation | Cholesterol-membran<br>e | Cholesterol-wate<br>r | Calcitriol-membran<br>e | Calcitriol-wate<br>r |
| --- | --- | --- | --- | --- |
| CMD_1CHOL | $0.5 \pm 0.6$ | $1.4 \pm 0.8$ | | |
| CMD_1CALC | | | $1.0 \pm 0.8$ | $1.8 \pm 1.2$ |
| CMD_16CHOL | $7.6 \pm 2.3$ | $22.4 \pm 3.4$ | | |
| CMD_16CALC | | | $14.9 \pm 2.9$ | $28.3 \pm 4.6$ |
| CMD_16CALC_16CHO<br>L | $7.7 \pm 2.4$ | $22.3 \pm 3.5$ | $14.6 \pm 3.0$ | $28.1 \pm 5.0$ |
| CMD_48CALC | | | $40.6 \pm 5.9$ | $84.6 \pm 7.3$ |
| CMD_48CALC_16CHO<br>L | $7.2 \pm 2.4$ | $21.5 \pm 3.7$ | $37.4 \pm 5.3$ | $87.4 \pm 8.3$ |

**Table 5.** Number of contacts between cholesterol-membrane, cholesterol-water, calcitriol-membrane, and calcitriol-water calculated in conventional molecular dynamics simulations.

| acronym | cholesterol-membrane | cholesterol-water | calcitriol-membrane | calcitriol-water |
| --- | --- | --- | --- | --- |
| CMD_1CHOL | $391 \pm 23$ | $9 \pm 5$ | | |
| CMD_1CALC | | | $393 \pm 29$ | $15 \pm 8$ |
| CMD_16CHOL | $4695 \pm 220$ | $135 \pm 19$ | | |
| CMD_16CALC | | | $4721 \pm 247$ | $228 \pm 33$ |
| CMD_16CALC_16CHOL | $4450 \pm 222$ | $137 \pm 19$ | $4825 \pm 168$ | $232 \pm 32$ |
| CMD_48CALC | | | $9402 \pm 380$ | $605 \pm 47$ |
| CMD_48CALC_16CHOL | $4193 \pm 209$ | $140 \pm 20$ | $8643 \pm 448$ | $613 \pm 49$ |

For simulations with 16 cholesterol molecules (CMD_16CHOL, CMD_16CALC_16CHOL, and CMD_48CALC_16CHOL), there was no significant difference in the number of contacts between cholesterol and water. Nonetheless, the number of contacts between cholesterol and the DMPE/DMPG membrane decreased as the number of calcitriol molecules increased. For simulations with 16 calcitriol molecules (CMD_16 CALC and CMD_16CALC_16CHOL), there was no significant difference in the number of contacts between calcitriol and water and between calcitriol and the membrane. However, the number of contacts between calcitriol and water was higher than that between cholesterol and water.

### 3.4 Water permeation through calcitriol-containing membranes

The average APL, sterol tilt angles, and water density at the bilayer center were higher, while D_P-P_ and S_CD_ were lower in calcitriol-containing membranes, indicating that these membranes were more disordered than cholesterol-containing membranes. Looser packing of lipids in the former facilitated water permeation.

The process of water permeation through the membrane in CMD_16CALC is shown in Figure 5. Only one water molecule crossed the membrane at 992-995 ns. The three-dimensional coordinates and Z distance of this molecule are shown in Figures 5A and B. The number of hydrogen bonds between this water molecule and calcitriol-containing membranes in different regions of the membrane is shown in Figure 5C. Upon crossing the membrane, this water molecule tended to form hydrogen bonds with hydrophilic groups, and upon entering the hydrophobic tail region, tended to form hydrogen bonds with calcitriol. Snapshots of the water permeation process are shown in Figures 5D-I.

**Figure 5.**
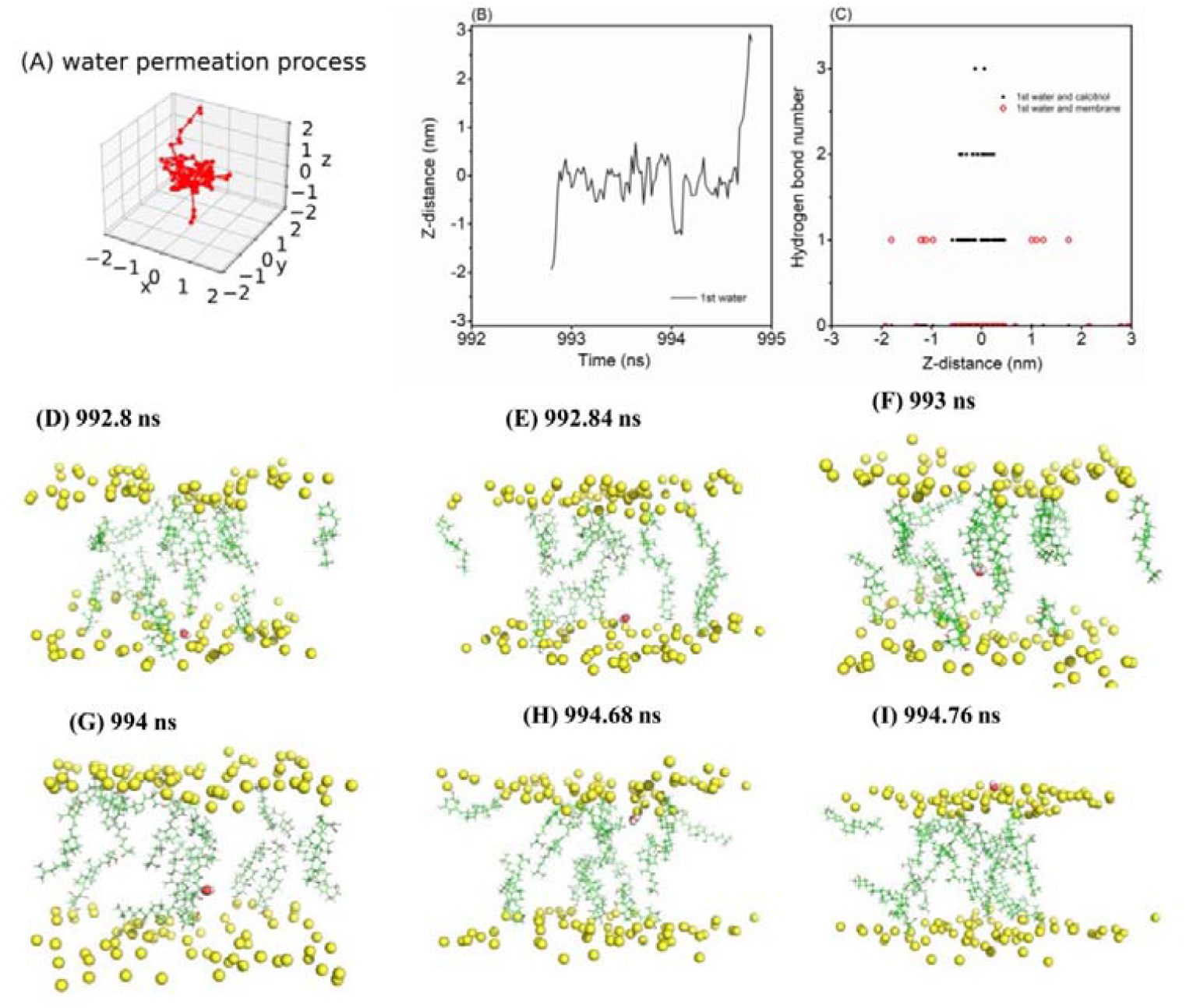
(A). The three-dimensional coordinate of a water molecule across the membrane bilayer. (B). Z distance between the water molecule and the membrane. (C) Scatter diagram of Z distance versus the number of hydrogen bonds. (D-I) The trajectory of a water molecule across the membrane bilayer. Hydrophilic heads and calcitriol molecules are shown in yellow and green. The oxygen and hydrogen atoms of water molecules are shown in red and white.

The permeation of three water molecules through the membrane in simulation CMD_48CALC is shown in Figure 6. These water molecules crossed the membrane at 956-960 ns. The three-dimensional coordinates and Z distances of these molecules are shown in Figures 6A-D. The number of hydrogen bonds between these molecules and calcitriol-containing membranes in different regions of the membrane is shown in Figure 6E. Upon crossing the membrane, these molecules tended to form hydrogen bonds with hydrophilic groups, and upon entering the hydrophobic tail region, tended to form hydrogen bonds with calcitriol. Snapshots of the water permeation process are shown in Figures 6F-I. The number of water molecules that crossed the membrane varied with the number of calcitriol molecules.

**Figure 6.**
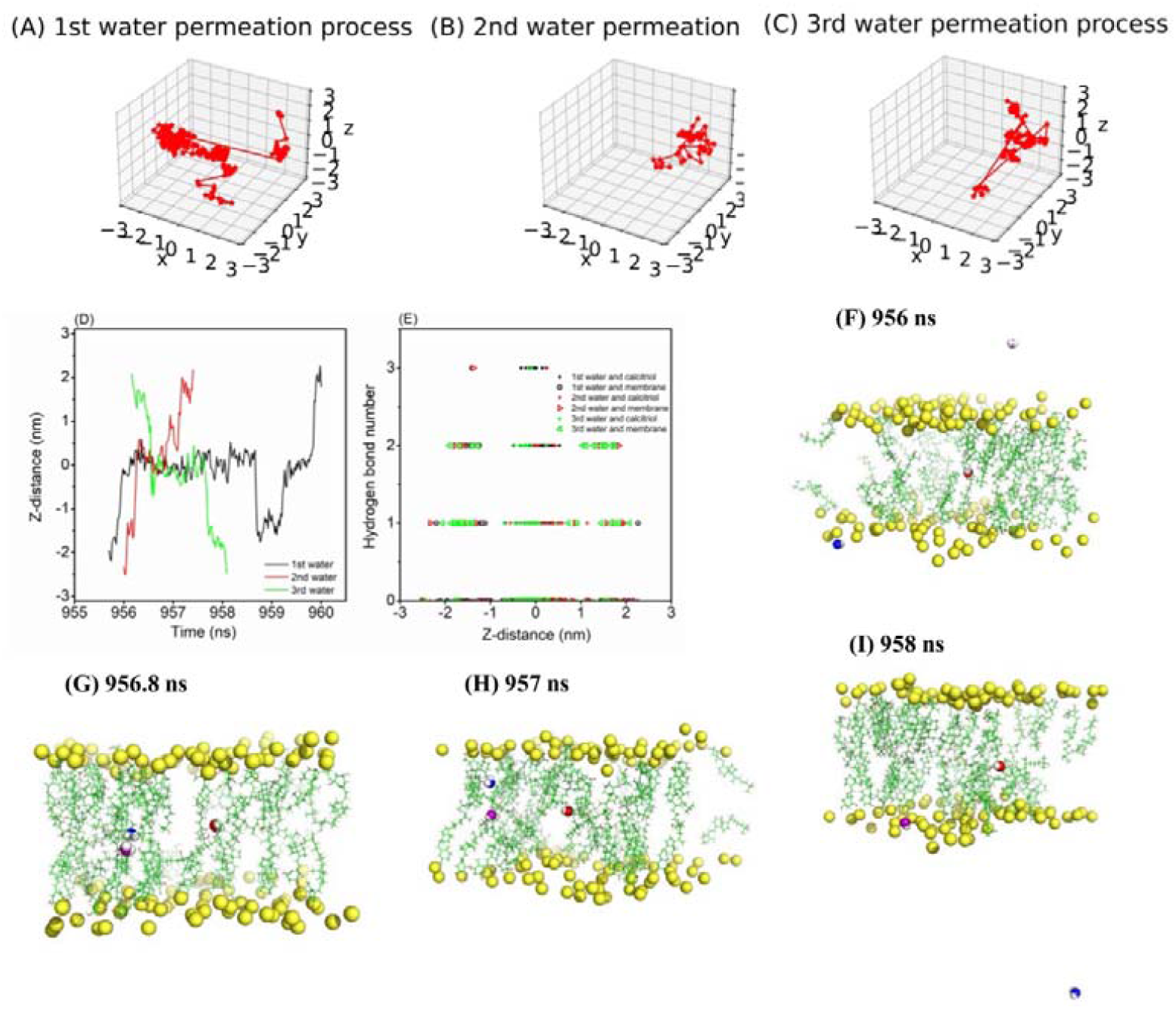
(A-C) The three-dimensional coordinates of three water molecules across the membrane bilayer. (D) Z distance between water molecules and the bilayer. (E) Scatter diagram of Z distance versus the number of hydrogen bonds. (F-I) The trajectory of water molecules across the bilayer. Hydrophilic heads and calcitriol molecules are shown in yellow and green. Water molecules are shown in red, purple, and blue.

### 3.5 Translocation of cholesterol and calcitriol from one layer to another

The translocation of cholesterol and calcitriol from one layer to another layer is essential for membrane structure, dynamics, and signaling. There were apparent differences in the two-dimensional free energy (ΔG) for the translocation of cholesterol and calcitriol into the DMPE/DMPG (3:1) bilayer (Figure 2). Calcitriol crossed from one layer to another more easily than cholesterol. The free energy barrier for the translocation of cholesterol (approximately 30 kJ/mol) was similar to the energy for translocating cholesterol in POPC and DPPC membranes[27]. The free energy barrier for the translocation of calcitriol was approximately 20 kJ/mol. This difference explains why calcitriol translocation was observed more frequently in CMD_16CALC than in CMD_16CHOL. The scatter plot of 16 cholesterols in CMD_16CHOL as CV1 and CV2 is shown in Figure S1. In the 1000 ns simulation, only one cholesterol molecule flip-flopped. The scatter plot of 16 calcitriol molecules in CMD_16CALC as CV1 and CV2 is shown in Figure S2. In the 1000 ns simulation, five calcitriol molecules were translocated. The time series of CV1/CV2 and tilt angles at 600-1000 ns for translocating five calcitriol molecules and one cholesterol molecule in CMD_16CALC and CMD_16CHOL are shown in Figure S3. Hence, the translocation of cholesterol and calcitriol across membrane leaflets was distinct. Snapshots of cholesterol and calcitriol translocation are shown in Figure 7. The flip-flop mechanism of cholesterol is similar between DMPE/DMPG, DPPC, POPC, DAPC membranes[27] and DPPC/DOPC membrane[28]. Cholesterol prefers to change orientation and then moves to the bilayer center, where sterol rings tilt approximately 90° with the membrane Z-axis. The tilt angle changes abruptly from ∼20° to ∼100° and the hydroxyl group locates from the membrane-water interface to the bilayer center. After crossing the bilayer center, cholesterol changes orientation again so that the hydroxyl group is positioned at the membrane-water interface. Unlike the flip-flop mechanism, calcitriol moves across the bilayer center vertically without changing its orientation. Translocation is similar to the bobbing mechanism of 27-hydroxycholesterol[29]. After crossing the bilayer center, the tilt angle changes from ∼30° to ∼100° so that the three hydroxyl groups are positioned at the membrane-water interface. Then the tilt angle changes again so that the 25-hydroxyl groups is positioned at the bilayer center.

**Figure 7.**
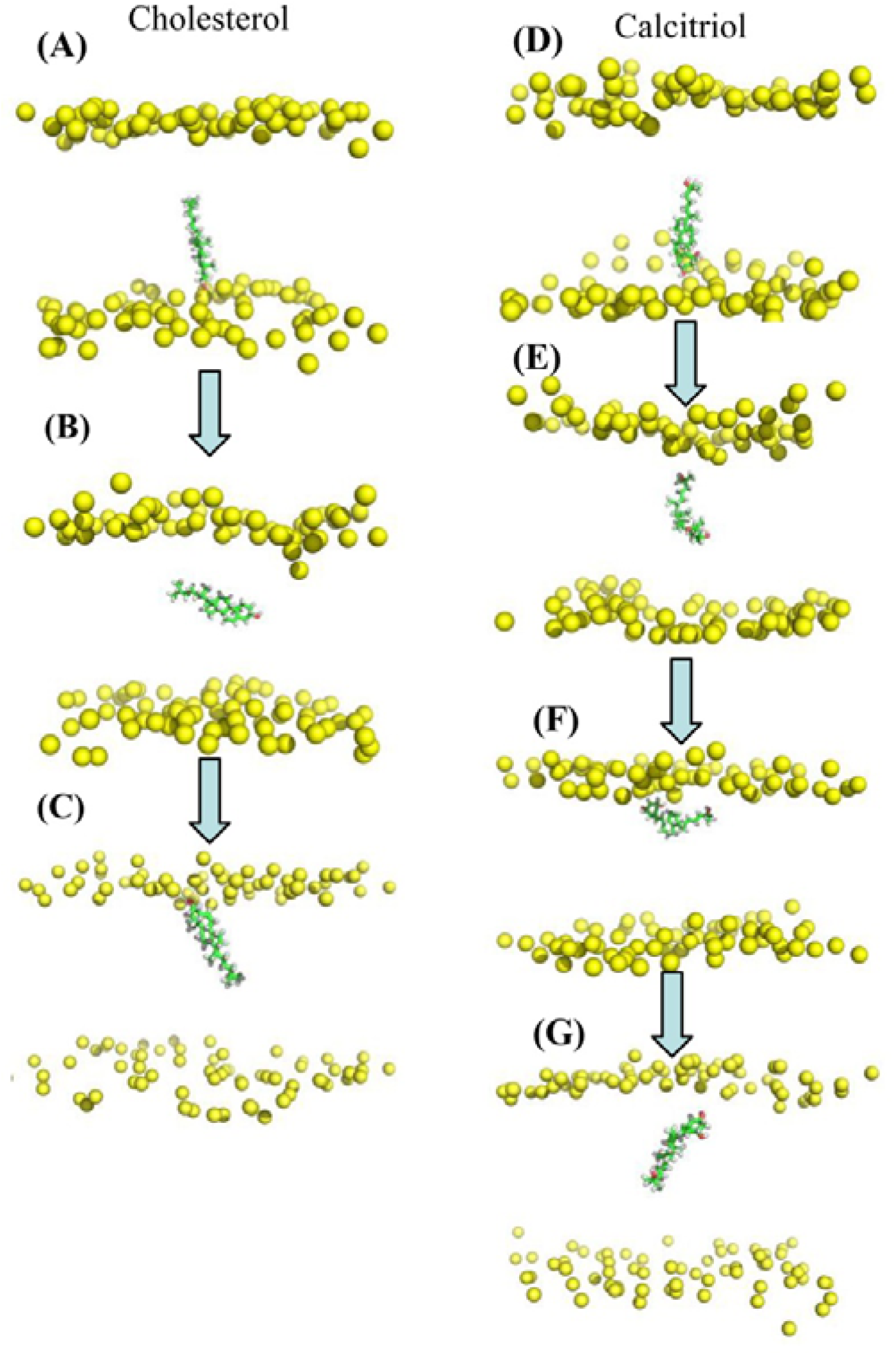
Snapshots of the translocation of (A-C) cholesterol and (D-G) calcitriol across the membrane. Hydrophilic heads and cholesterol/calcitriol molecules are shown in yellow and green. The oxygen atoms of hydroxyl groups are shown in red.

## 4. DISCUSSION

The chemical structure of calcitriol and cholesterol is similar. Calcitriol and cholesterol affect the physical properties of lipid bilayers by increasing S_CD_ and membrane thickness. However, cholesterol increases these parameters more strongly than calcitriol. Our results indicate that cholesterol-containing membranes are more ordered than calcitriol-containing membranes. Our simulations showed that the average APL and sterol tilt angles were slightly larger in calcitriol-rich membranes. Looser packing of lipids in these membranes facilitated calcitriol translocation. Moreover, calcitriol increased water permeability across the membrane, while cholesterol had the opposite effect.

Oxidized oxysterols such as 27-hydroxycholesterol and 25-hydroxycholesterol increase membrane permeability. Baker et al.[30, 31] showed that cholesterol and 25-hydroxycholesterol had different effects on POPC bilayers, which might be due to the wider range of orientations of the latter. Olzynska and Kulig et al.[29, 32] compared the effects of 7β-hydroxycholesterol and 27-hydroxycholesterol and found that the latter facilitated water transport across the membrane; this process was also associated with the local deformation of the lipid bilayer (bobbing).

The long axis of cholesterol was positioned at an angle of approximately 20° to the membrane axis, and the tail extended into the bilayer center. Calcitriol assumed several orientations but did not align perfectly with the bilayer. The long axis of calcitriol was positioned at an angle of approximately 30° to the membrane axis. Nonetheless, some calcitriol molecules were positioned parallel to the membrane surface, maintaining both hydroxyl groups away from the hydrophobic core.

Calcitriol crosses each membrane leaflet more easily than cholesterol. Moreover, the translocation path of these two molecules is quite different (Figure 7). Cholesterol molecules are positioned parallel to the membrane surface and flip-flop at the bilayer center. In turn, calcitriol moves across the bilayer center without changing its orientation along the membrane Z-axis, becomes parallel to the membrane surface at the membrane-water interface, and then rotates approximately 90º at this interface.

## 5. CONCLUSIONS

We demonstrated that calcitriol improved water permeability across the DMPE/DMPG (3:1) lipid bilayer, while cholesterol had the opposite effect. Previous studies[33, 34] and our results suggest that cholesterol decreases water permeability in several bilayer systems, including DPPC, DOPC, SOPC, and DMPE/DMPG. Moreover, the hydroxylation of the hydrocarbon tail of sterols increases the translocation of molecules across the membrane, in line with a previous study[27]. Molecular dynamics simulations suggest that the translocation mechanism and free energy barrier of calcitriol and cholesterol differ, and deformations in the bilayer structure increase water permeability. These results corroborate the distinctive role of calcitriol in *H. pylori* infection.

## Acknowledgments

We acknowledge the support from the National Natural Science Foundation of China (NSFC, grant numbers 32171249 and 31670727), Youth Innovation Team Lead-education Project of Shandong Educational Committee. We are grateful to TopEdit (www.topeditsci.com) for language editing.

## Data Availability

The data that support the findings of this study are available from the corresponding author upon reasonable request.

## Supplementary Material

Figures S1-S3.

## Author Contributions

Zanxia Cao, Liling Zhao, Mingcui Chen and Lei Liu performed MD simulations, Zanxia Cao, Liling Zhao and Lei Liu analyzed the data, Zanxia Cao and Lei Liu drafted up and revised this manuscript.

